# The distribution of the time to the most recent pairwise genetic common ancestor of a set of sampled genomes

**DOI:** 10.1101/2025.07.16.665208

**Authors:** Zehui Zhao, Rohan S Mehta, Daniel B Weissman

## Abstract

In a sample of chromosomes from a recombining population, each pair of individuals will have different most recent common genetic ancestors at different loci. We consider the distribution of the time to the most recent of these most recent common ancestors—the most recent time at which any pair of individuals in the sample share a common genetic ancestor at any locus. We use simple heuristic arguments, formal calculations, and coalescent simulations to find that as long as the chromosomal map length *R* is sufficiently long and the sample size *n* is not too large, the distribution of this time is peaked around a characteristic value. This value has the unusual scaling 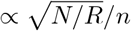, where *N* is the effective size of the population.

## Introduction

In a recombining population, each region of the genome will have its own set of ancestral relationships among individuals, i.e., its own ancestral tree. This sequence of trees across the genome is the mediator connecting population genetic data and historical population dynamics. The large number of trees across the genome means that even a single diploid individual or a small sample of individuals contains a wealth of information about the historical dynamics of the population [e.g. Li and Durbin, 2011, Weissman and Hallatschek, 2017]. But taking advantage of this information requires understanding the distribution of trees produced by different demographic histories. One major challenge is that the space of possible trees relating large samples of individuals is very high dimensional and will only be sparsely sampled even if genomes are long. Many methods therefore rely on lower-dimensional summary statistics—specific features of trees. For instance, one can consider the distribution of times to the most recent common (genetic) ancestor (TMRCAs) of all pairs of sampled individuals. The minimum of this distribution of TMRCAs (i.e., the time to the most recent most recent common ancestor of a pair) plays a particularly important role in the powerful MSMC method for demographic inference [Schiffels and Durbin, 2014], as well as the new IICR_*k*_ method [Chikhi et al., 2024]. However, to our knowledge, there is no known expression for the expected distribution of the minimum TMRCA in even the simplest demographic scenario of a constant, well-mixed population.

Across the genome, coalescence times can span a wide range. At a random gene, the time for two random individuals to share a common ancestor (coalesce) will typically be on the order of the effective population size; for humans, this is roughly 10,000 generations [Kruglyak, 1999]. On the other hand, the minimum possible coalescence time for those two individuals is the time till they first share a genealogical common ancestor, which often grows only logarithmically with population size. For humans, this time is about 20 generations [Chang, 1999], i.e., two orders of magnitude smaller than the typical coalescence time. Intuitively, the difference between these timescales arise because genes have exactly one ancestor in the previous generation, and thus the total number of ancestral lineages is constant or decreasing going back in time, whereas individuals have two, and so the total number of ancestors grows approximately exponentially [Chang, 1999, Derrida and Jung-Muller, 1999]. When we consider the TMRCA of a sample of individuals with recombination, the situation is intermediate between the two extremes of single-gene ancestry and pedigree ancestry. The number of ancestors does increase going back in time because of recombination, but not all genealogical ancestors are genetic ancestors. Even among past individuals who are genetic ancestors of two samples, many will not be ancestral at the same region of the genome, and will therefore not be genetic common ancestors.

The general solution for the ancestry of a set of individuals with recombination is the Ancestral Recombination Graph (ARG) [Griffiths and Marjoram, 1996], which describes how ancestral lineages are broken up by recombination in addition to coalescing. The ARG has a notoriously large state space and it is therefore usually extremely computationally intensive to deal with it in full, although the recent developments are promising [e.g. Kelleher et al., 2016, 2019]. Several methods have been developed to approximate the ARG to make computations simpler [e.g. Rasmussen et al., 2014, Kelleher et al., 2019, Speidel et al., 2019, Schaefer et al., 2021, Zhang et al., 2023]. One such method was developed by McVean and Cardin [2005] and is called the Sequentially Markovian Coalescent (SMC). This method only allows lineages to coalesce if they are genetic ancestors of overlapping regions of the genome in the sampled individuals. A slight modification of the SMC, the SMC^*′*^, also allows coalescence between lineages that are ancestral to adjacent regions of genome, rather than just overlapping regions [Marjoram and Wall, 2006]. Both the SMC and SMC^*′*^ are approximations to the ARG, with the SMC^*′*^ being more accurate [Wilton et al., 2015]. One notable application is in demographic inferences, with the SMC underlying PSMC [Li and Durbin, 2011] and the SMC^*′*^ underlying MSMC Schiffels and Durbin [2014].

In this paper, based on approximations equivalent to SMC and SMC^*′*^, we find approximate expressions for the distribution of the genome-wide TMRCA of a sample of haploid individuals and verify their accuracy with simulations of the full ARG. We find that the expected genomewide TMRCA scales as the square root of the population size divided by the total genetic map length of the sequenced region 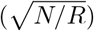. This dependence on population size is intermediate between the linear dependence of the single-gene TMRCA and the logarithmic dependence of the genealogical TMRCA, while the dependence on the map length is a new feature not present in these two other TMRCAs. For very large sample sizes and limited recombination, there is a transition to a different scaling behavior, where the most closely related pair of individuals coalesced so recently that recombination had no time to act, and the genome-wide TMRCA reduces to the single-gene TMRCA. All code used for simulations or plotting in this paper can be found in https://github.com/weissmanlab/Recombination_MRCA.

## Model and background

We consider a population of *N* haploid individuals with random mating. Equivalently, this could be seen as a population of *N/*2 diploid individuals. Let *τ* be the time to the present measured in units of *N* generations, Each individual’s genome consists of a single chromosome of map length *R*. It will be convenient to use time *τ* measured in units of *N* generations, so that pairs of lineages coalesce at rate one. In terms of this scaled time, our results will depend on the parameters *N* and *R* only through the scaled map length *ρ* ≡ *NR*. To convert them to unscaled time *t* in units of generations, we simply need to multiply by *N* : *t* = *Nτ*.

We wish to find an expression for the probability density *p*(*τ*) of the time *𝒯* to the most recent common genetic ancestor of any pair of individuals in a sample of size *n*. The mean and standard deviation *𝒯* of are useful ways of summarizing this distribution; we will denote them by 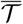 and *σ*, respectively. We will refer to *p*(*t*) as the “TMRCA distribution”. Equivalently, we can consider the probability *F* (*τ*) that no common genetic ancestors have occurred by time *τ*. (*F* is called the survival function or the complementary cumulative distribution.) Both of these can be written in terms of the rate *c*(*τ*) at which common genetic ancestors occur at time *τ* given that none have occurred up to that time:

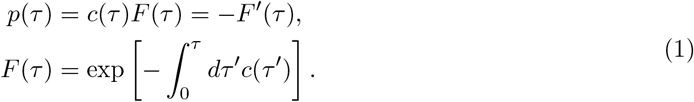

The problem is thus reduces to finding *c*(*τ*) as a function of scaled map length *ρ* and sample size *n*. For a single locus, i.e., in the absence of recombination (*ρ* = 0), until the first coalescence at time *𝒯* there are always exactly *n* lineages. A coalescence between any pair of lineages would result in a genetic common ancestor, so the total rate is just the number of pairs:

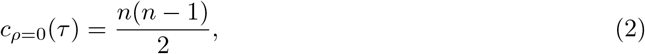

and the TMRCA distribution is simply exponential:

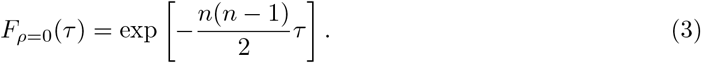

The mean and standard deviation for the TMRCA are:

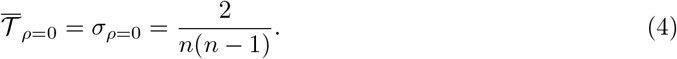

Intuitively, the typical TMRCA is *𝒪* (1*/n*^2^)(linear in the population size in unscaled time) and is moderately broadly distributed, with standard deviation equal to its mean. Note that for a sample of just *n* = 2 individuals, *n*(*n −*1)*/*2 = 1. So *F*_*ρ*=0_(*τ*) = [*F*_*ρ*=0,pairwise_(*τ*)]^*n*(*n−*1)*/*2^, i.e., coalescence is independent between sampled pairs.

Recombination causes the number of lineages to increase stochastically as we go back in time (Fig. 1). This greatly increases the number of pairs of lineages, but now only a minority of pairs of lineages are ancestral to overlapping regions of the sampled genome. It is only this minority of pairs whose coalescence causes a genetic common ancestor. To emphasize the importance of events involving this minority of pairs, Kelleher et al. [2016] reserve the term “coalescence” for these events that lead to a *genetic* common ancestor and use “common ancestor event” to generically refer to both these and the more common case in which the two merging lineages are ancestral to disjoint regions of the genome. For the rest of this paper, we will follow this convention. *c*(*τ*) is thus the rate of *coalescence* at time *τ*, conditioned on no coalescence having already occurred.

**Figure 1:**
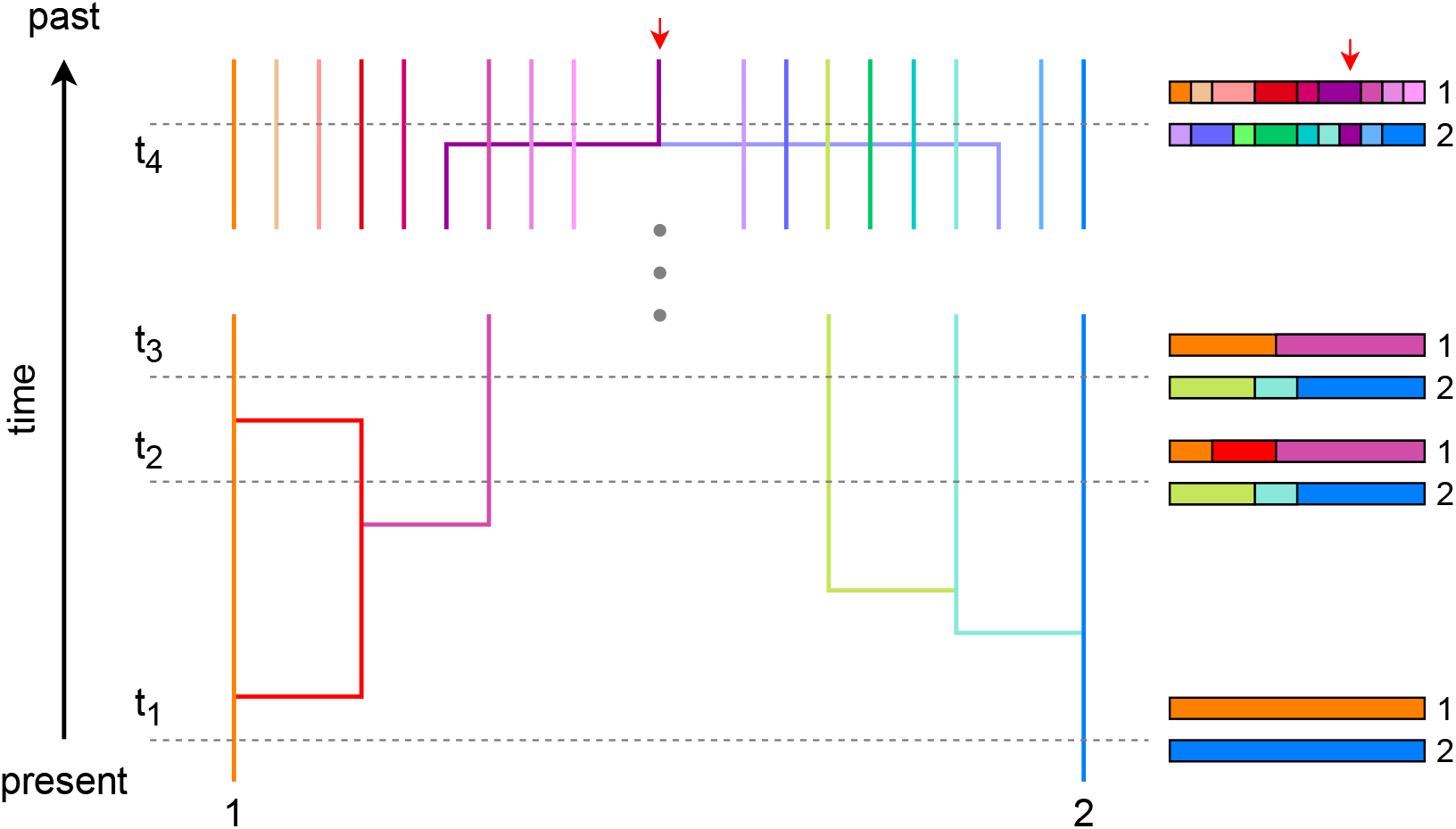
Schematic of the ancestral recombination graph of a sample of *n* = 2 genomes up to the time of the most recent common genetic ancestor. To the right are shown the regions of the sampled genomes that each lineage is ancestral to at each of the labeled times. Immediately after sampling, at *t*_1_, there is just one lineage ancestral to the whole genome of each sample. Moving backward in time, recombination breaks these up into multiple lineages, each ancestral to only part of the genome. Lineages that are ancestral to different parts of the genome can come together in “common ancestor events” (orange and red merger between *t*_2_ and *t*_3_) which do not create a common genetic ancestor [Kelleher et al., 2016]. The first coalescence between lineages ancestral to overlapping regions of the genome is between the orange and blue lineages just prior to *t*_4_.

## Analysis and results

### Heuristic analysis

We begin with a back-of-the-envelope asymptotic analysis to guide intuition. Above, in the asexual case, we saw that the typical TMRCA was *τ*_*ρ*=0_ ∼ 2*/*(*n*(*n* −1)). For recombination to greatly reduce this, it must occur often during this time, *ρ* ≫ *n*(*n* − 1). Once recombination has had a chance to act, *ρτ* ≫1, the ancestry of each sampled genome will be split into ∼ *ρτ* blocks. Each block overlaps with ∼ 2(*n* − 1) blocks from the ancestry of the other *n* − 1 samples. (The factor of two is because the recombination breakpoints occur at different places in different lineages, so each block on average overlaps with two other per other sampled individual.) So there are a total of *c*(*τ*) ∼ *n*(*n* − 1)*ρτ* pairs of independently coalescing blocks. Combining this with Eq. 1 and Eq. 3, we have asymptotic expressions for the probability of the TMRCA being older than *τ* :

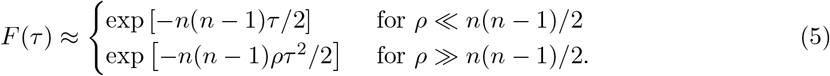

In the first case, limited recombination or very large sample size, the first coalescence will usually occur before the first recombination. In the second case, the genomes typically recombine many times before the first coalescence. Inspecting the second line of Eq. 5, we see that the typical TMRCA in this case scales as 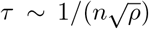 (neglecting constant factors), or 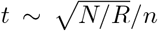 in unscaled time.

While we will see below that this intuitive argument is essentially correct, it only gives us asymptotic expressions. And even these are limited: in particular, the second line of Eq. 5 must only apply for intermediate values of time *τ*, near the typical TMRCA. It must break down for very recent times, *τ* → 0, as it predicts that the coalescence rate *c* drops to zero, when in fact it must be bounded below by the asexual value *n*(*n* − 1)*/*2. This is straightforward to fix, but a more difficult issue is that it must also break down for very ancient times, *τ* → ∞, as it predicts that the coalescence rate *c* grows without bound, giving a Gaussian tail for *F*. In fact, the tail must be no steeper than exponential. To see this, note that *F* is bounded below by the probability of never having a recombination event and none of the *n* lineages coalescing, exp [− *nρτ* − *n*(*n* − 1)*τ/*2]. Below, we will use the SMC and SMC^*′*^ to derive improved versions of Eq. 5 that correct these issues.

### The rate of coalescence conditional on the number of ancestral lineages

Let *𝒜* (*τ*) denote the ARG up to time *τ*, a random variable. In the single-locus case above, the TMRCA distribution was easily derived because until the first coalescent event, *𝒜*always consisted of exactly *n* lineages that were all ancestral to the same genetic region. But for multiple loci, the stochasticity in recombination and non-coalescent common ancestor events mean that each realization will have a different ARG and therefore a different instantaneous coalescence rate. For example, a sample whose history involves an unusually high number of recombination events will have an unusually high number of genetic ancestors, and therefore an unusually high coalescence rate. So in general, finding the TMRCA distribution requires finding the coalescence rate conditional on the ARG, *c*(*τ* |*𝒜* (*τ*)), and averaging this over the distribution of ARGs. However, even parameterizing the space of ARGs is hard because it is so high-dimensional.

To simplify the analysis, we make what is essentially the same approximation as the SMC^*′*^: that each lineage is ancestral to the sample at just one connected region of its genome. Intuitively, this assumption reflects the fact that any associations between ancestry at distant regions of the genome are likely to be fleeting as they will quickly be broken up by recombination. We will now show that under this SMC^*′*^ approximation the conditional rate of coalescence *c*(*τ* | *𝒜* (*τ*)) depends on the ARG *𝒜* (*τ*) only through the total number of lineages at time *τ*. We will do this by calculating the instantaneous coalescence rate *č* by induction on the total number of lineages. Let this total number of lineages be *n* + *X*(*τ*). The random variable *X* tracks how many extra lineages have been created by recombination, minus those that have been lost to non-genetic common ancestor events. For example, in Fig. 1, we start with a sample of *n* = 2 haplotypes, and at time *t*_3_, after four recombination events and one common ancestor event, we have *X* = 4 − 1 = 3 and the total number of lineages is *n* + *X* = 5. Given that we are only tracking the dynamics until the first coalescence event, *X* must always be non-negative. Our claim is that there is that we can write *c*(*τ* | *𝒜* (*τ*)) = *č*(*X*(*τ*)) for some function *č*.

For the base case of the induction, *X* = 0, i.e., *n* lineages in total where each sample has exactly one ancestor for its whole genome, we simply have the single-locus expression *č*(0) = *n*(*n* −1)*/*2. For the induction step, we will find how *č* increases when *X* increases by one. Recall that *X* increases by one when a recombination event breaks the ancestral segment in a lineage into two lineages, each with one ancestral segment. Because no genetic coalescence has yet occurred, the broken segment is ancestral to exactly one of the *n* sampled individuals. Each of the other *n* − 1 individuals has a unique ancestor carrying the region of the genome containing the breakpoint. These *n* − 1 ancestral lineages now have an additional lineage that they can each coalesce with, so the number of pairs *č* has increased by *n* − 1. For example, in Fig. 2, we start with a possible ARG for a sample of *n* = 3 genomes on the left, where a recombination event occurs on the yellow lineage and changes the ARG to the right. Before the recombination, there are a total of *č* = 5 independently coalescing pairs, and after the recombination, there are *č* = 7, i.e., the number of pairs *č* has increased by *n* − 1 = 2. Here, we have assumed that no other individual’s ancestry has a recombination breakpoint in exactly the same spot, which will almost always be true for uniformly distributed breakpoints in a long genome. In the Appendix, we simulate a genomic region of only 100 loci, violating this assumption, and verify that this has only a small effect on the TMRCA distribution. Given the starting value *č*(0) and the fact that it increases by *n* − 1 for each increment in *X*, we obtain:

**Figure 2:**
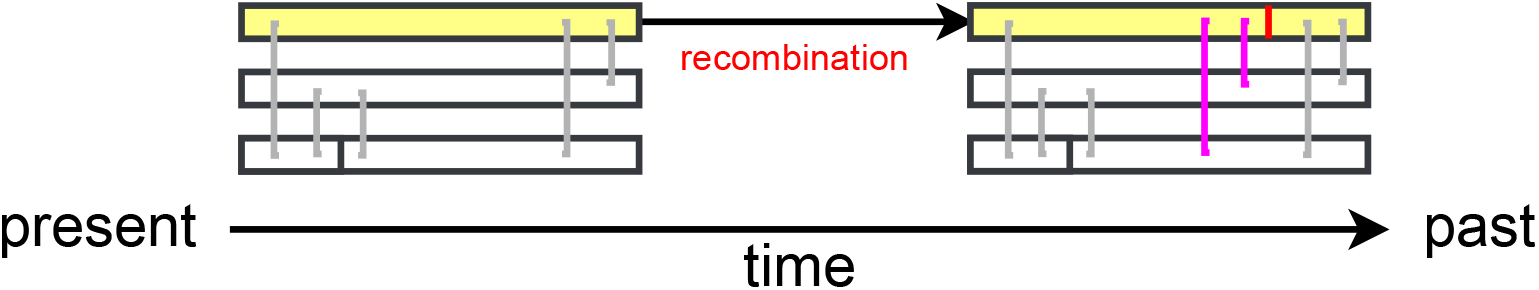
Schematic of recombination affecting the coalescent rate in a sample of *n* = 3 genomes. Closer to the present (left), there are four lineages (because the bottom lineage has undergone a recombination event, creating *X* = 1 excess lineage) and *č* = 5 independently coalescing pairs, shown in gray. Deeper in the past, a recombination event (vertical red line) occurs in the ancestry of the highlighted top genome, splitting the lineage into two, creating *n* − 1 = 2 new pairs of independently coalescing pairs, shown in pink, for a total of *X* = 2 excess lineages and *č* = 7 independently coalescing pairs.

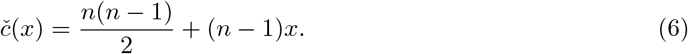

For example, at time *t*_3_ in Fig. 1, *X*(*t*_3_) = 3 excess lineages and *č*(3) = 2(2 −1)*/*2 + 3(1) = 4 pairs that are ancestral to overlapping genome segments and can therefore coalesce (orange and green, orange and cyan, purple and cyan, and purple and blue). It may seem surprising that the coalescence rate *č* only grows linearly with the excess number of lineages *X*, given that *č* is quadratic in the number of starting lineages *n*. Indeed, the total rate of common ancestor events *is* quadratic in *X*; it is just that coalescent events make up an increasingly small fraction of these as the genome gets more split up by recombination.

Technically above we have only shown that Eq. 6 is true under the SMC approximation that neglects all non-genetic common ancestor events, so that *X* only increases until the time *𝒯* of the TMRCA. Under the more accurate SMC^*′*^ approximation, we also allow non-genetic common ancestor events, but only of adjacent blocks of the genome. Each of these events just reverses the splitting due to recombination considered above, and so just reduces *X* by one and *č* by *n* − 1, as required for Eq. 6 to be true. To see why it is important to neglect non-genetic common ancestor events involving non-adjacent genome blocks, consider what would have happened in Fig. 1 if instead of the orange and red lineages rejoining at time *t*_3_ it had been the orange and purple lineages. Then *X* would still have decreased to three, but there would be five possible coalescent pairs (pink/orange with blue, cyan, or purple, and yellow with blue or cyan).

Given that the instantaneous coalescence rate depends only on *X*, we can rewrite Eq. 1 as:

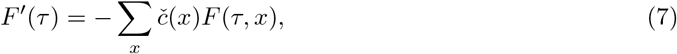

where *F* (*τ, x*) is the joint probability of not yet coalescing by time *τ* and having *X*(*τ*) = *x* excess lineages (so *F* (*τ*) = ∑_*x*_ *F* (*τ, x*)). To solve this, we need the full master equation for *F* (*τ, x*). In solving the master equation, it will be helpful to consider the generating function *ψ* defined by:

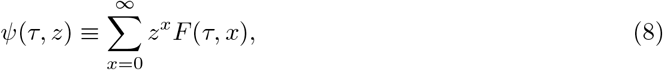

from which we can recover the survival function *F* (*τ*) via *F* (*τ*) = *ψ*(*τ*, 1). We obtain two sets of slightly different results under the SMC and SMC^*′*^ below. Within each section, we derive the corresponding TMRCA distribution and then compare it to the distribution obtained through full ARG simulations in msprime [Kelleher et al., 2016] to evaluate its accuracy.

### TMRCA distribution under the SMC approximation

We first consider the SMC approximation, which ignores non-coalescent common ancestor events entirely. In this case, the number of excess lineages *X*(*τ*) is just a Poisson process increasing at the recombination rate *nρ* (until we reach the first coalescent event and the process ends). The master equation is therefore:

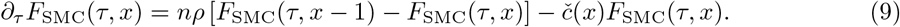

The first term on the right-hand side describes recombination incrementing the lineage number, and the second term on the right describes coalescence with the rate depending on the lineage number. (Recall that there is no chance of having negative excess lineages: *F* (*τ, x*) ≡ 0 for *x <* 0.)

Rewriting Eq. 9 in terms of the generating function *ψ* gives the partial differential equation:

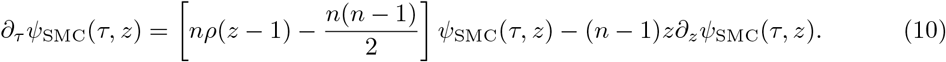

To obtain the boundary condition, note that there are no excess lineages at *τ* = 0 and no realization has coalescenced yet, so *ψ*(*τ* = 0, *z*) = 1. Eq. 10 can be solved using the method of characteristics to obtain:

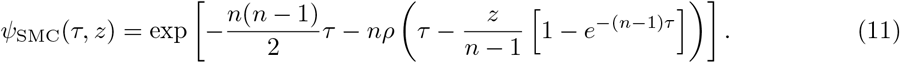

Evaluating Eq. 11 at *z* = 1 gives the marginal survival function:

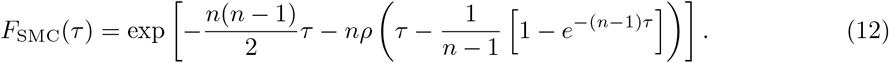

Eq. 12 simplifies in the short- and long-time limits:

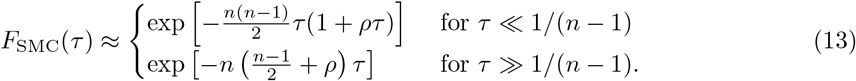

The probability density follows immediately from Eq. 12 as *p*_SMC_(*τ*) = −*F*_SMC_*′* (*τ*). Fig. 3 shows that *p*_SMC_ for the most part accurately matches the results of full coalescent simulations over a wide range of sample sizes and recombination rates. The main inaccuracy is that Eq. 12 overestimates the peak of *p* for small sample sizes and low recombination rates (upper left panel), and correspondingly underestimates the tail (inset).

**Figure 3:**
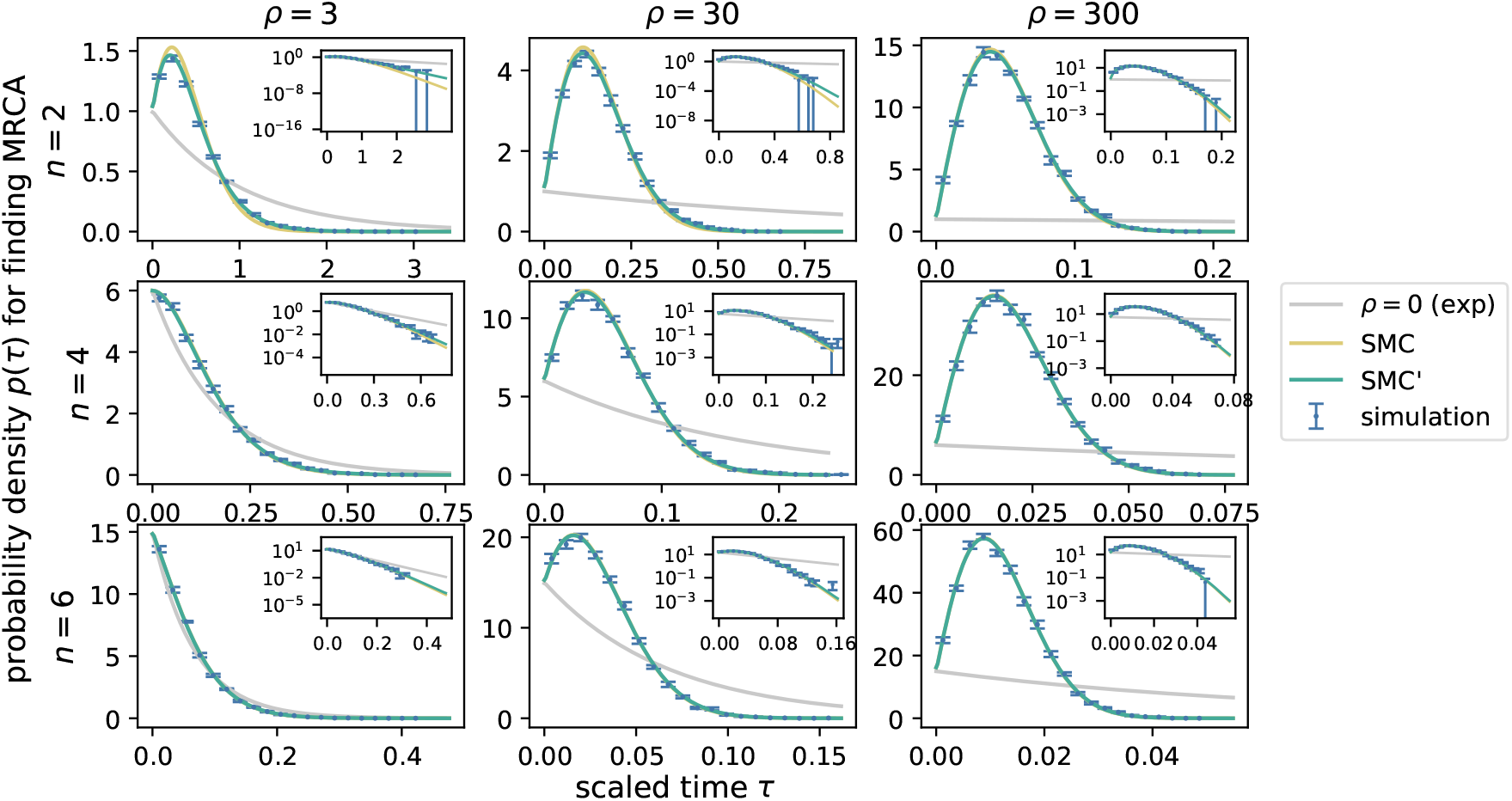
The TMRCA distribution is exponential for low recombination rate *ρ* ≪ *n*^2^ and peaked for high recombination rate *ρ* ≳ *n*^2^. Panels show the density of coalescence times *p*(*τ*) for different values of the sample size *n* and recombination rate *ρ*. The tail of the distribution is shown in log-log insets. For each *n* and *ρ*, points with error bars show the mean and standard error across 10 empirical distributions, each based on 1000 msprime tree sequence simulations. The gray curves show the exponential curve expected in the absence of recombination (derived from Eq. 3), and the gold and cyan curves show the densities corresponding to analytical distributions obtained under the SMC (Eq. 12) and SMC^*′*^ (Eq. 17), respectively. Both approximations match the simulated results well, with the SMC^*′*^ having higher accuracy than the SMC for small sample sizes and low recombination rates. Error bars represent standard errors in values.

Comparing the SMC expressions for the survival function Eq. 12 and Eq. 13 to the simple heuristic one Eq. 5, we see that the heuristic equation is just a further simplification of the short-time expression (first line of Eq. 13) in the two limiting cases *ρ* ≪ *n*^2^ and *ρ* ≫ *n*^2^. The SMC longtime behavior, however, captures a qualitative feature that we noted was missed by the heuristic argument: the distribution has an exponential tail rather than a Gaussian-like one. In terms of the coalescence rate *c*, this means that the SMC predicts that *c* saturates at a constant value, rather than continuing to increase linearly with time forever. In fact, the SMC prediction exactly matches our simple lower bound set by the probability of avoiding *both* recombination and coalescence. The insets in Fig. 3 confirm that the simulated distribution does indeed have an exponential tail, although with a slightly different rate constant *c* from that predicted by the SMC, particularly for small sample sizes and low recombination rates (top left).

### TMRCA distribution under the SMC′ approximation

While the distribution obtained under the SMC approximation is highly accurate, it still underestimates the TMRCA by having a higher than simulated peak and shorter tail. This is not surprising since SMC ignores all non-coalescent common ancestor events, and these ignored events reduce the number of lineages and thereby reduce the coalescent rate. To correct for this underestimation, we now consider the SMC^*′*^ approximation, which extends SMC by including non-coalescent common ancestor events that do not break the connected ancestral region approximation. More concretely, SMC^*′*^ allows lineages with adjacent ancestral segments to rejoin. An example is the common ancestor event just before *t*_3_ in Fig. 1 that rejoins the red lineage to the orange one.

To find the rate of adjacent lineages rejoining, note that the number of adjacent lineage pairs is the same as the excess lineage number *X*: every segment of the genome has an adjacent segment to its right, except for the *n* segments containing the right ends of the genomes which we can treat as the *n* “original” lineages. The rate of common ancestor events between adjacent lineages is therefore *X*. For example, at time *t*_3_ in Fig. 1, there are *X* = 3 excess lineages and three adjacent lineage pairs that could have a common ancestor event (pink-orange, blue-cyan, and cyan-purple).

The master equation for *F*_SMC_*′* (*τ, x*) is:

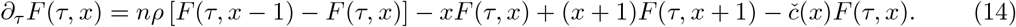

This is the same as Eq. 9 for *∂*_*τ*_ *F*_SMC_(*τ, x*), but with the additional middle terms accounting for common-ancestor events between adjacent lineages. The corresponding generating function *ψ*_SMC_*′* (*τ, z*) ≡ ∑_*x*_ *z*^*x*^*F*_SMC_*′* (*τ, x*) satisfies:

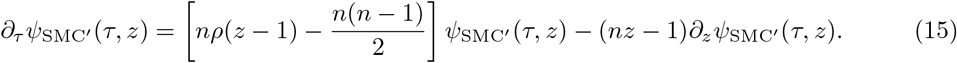

The boundary condition remains *ψ*(0, *z*) = 1.

This can again be solved by the method of characteristics to find:

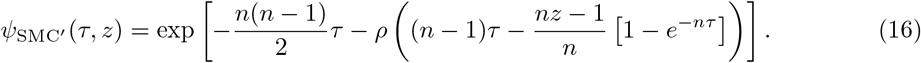

Evaluating at *z* = 1 gives the survival function:

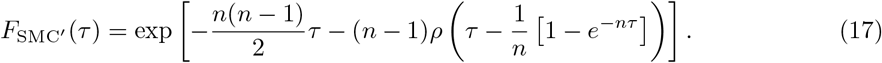

The TMRCA distribution 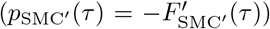 obtained under the SMC^*′*^ is shown in Fig. 3. As expected, it matches the simulated results even better than the SMC does, particularly at small sample sizes and modest recombination rates (upper left panel).

Eq. 17 has the short- and long-time limits:

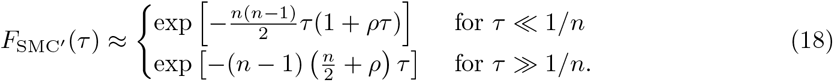

Comparing these to the SMC results Eq. 13, we see that the short-time limit is the same, while the long-time limit has slightly different dependence on sample size. This makes sense, as it is at the long times that the adjacent-lineage common ancestor events that differentiate the SMC and SMC^*′*^ will have had time to occur. As with the SMC, the form of the tail for the SMC^*′*^ has a simple interpretation: instead of being the probability of *completely* avoiding recombination (along with coalescence), it allows the fraction 1*/n* of recombination events where the new lineage rejoins with the lineage it broke off from in a common ancestor event instead of coalescing with any of the other lineages. The SMC^*′*^’s approximation for the long-time behavior is a better match to the simulations than the SMC’s (Fig. 3, insets, particularly *n* = 2 and *ρ* = 3).

### Mean and standard deviation of the TMRCA

The mean and standard deviation of the TMRCA under the SMC and SMC^*′*^ can be found from the survival functions Eq. 12 and Eq. 17 using numerical integration or in terms of special functions. The results closely match simulations (Fig. 4), as expected given the close match in the full distributions. The SMC^*′*^ is consistently *<* 5% from the simulated values. We also look for simple asymptotic approximations that are easier to interpret and allow us to understand the scaling with the parameters. One limit is simple: for limited recombination or large sample sizes (*ρ* ≪ *n*^2^), the results should reduce to the *ρ* = 0 case, with the mean and standard deviation approaching Eq. 4. This can be seen on the left-hand side of the bottom row of plots in Fig. 4.

**Figure 4:**
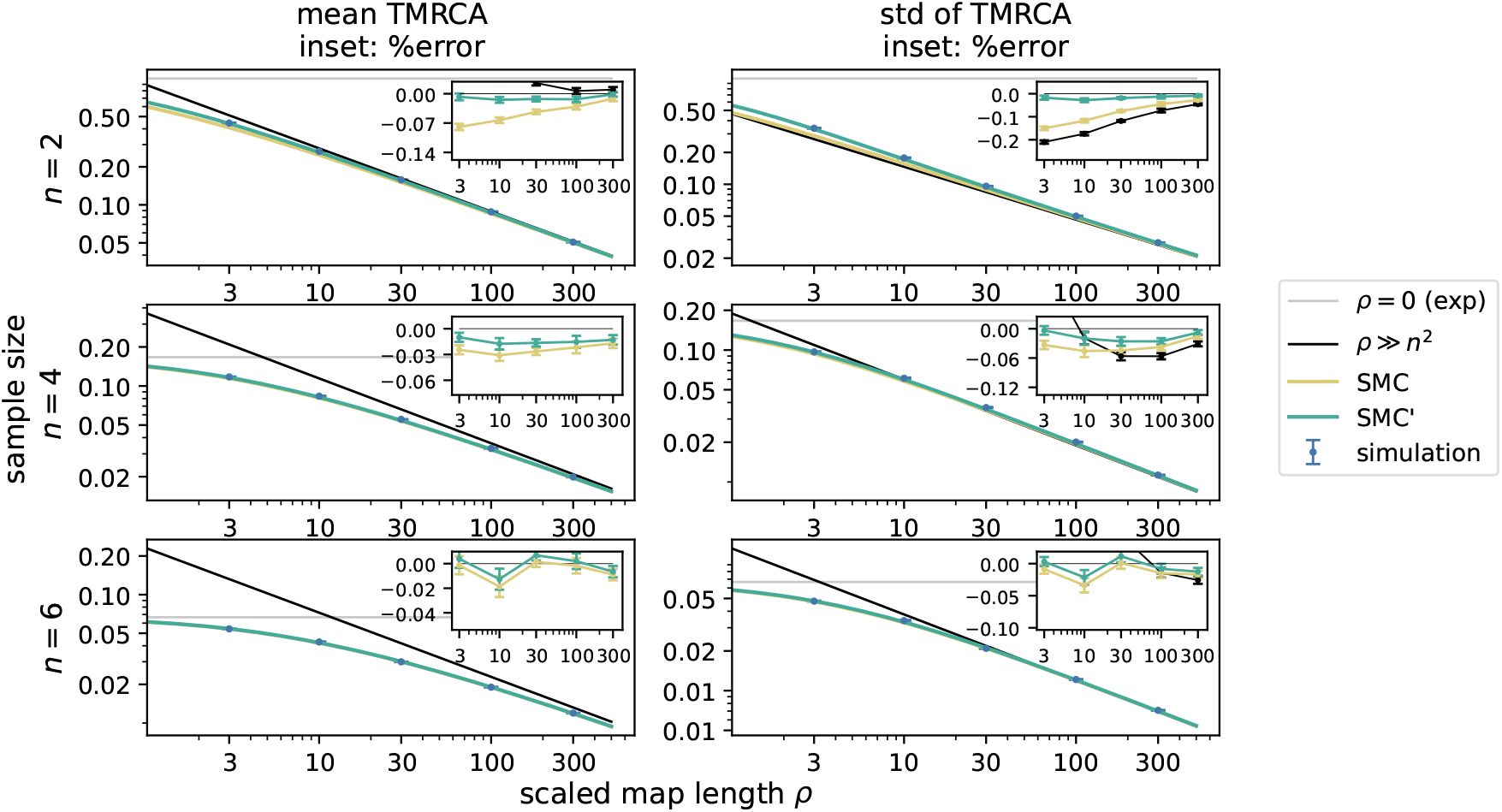
The mean and standard deviation of the TMRCA scale with the square root of the population size. Plots show the mean (left) and standard deviation (right) of *𝒯* for a range of values of the sample size *n* and recombination rate *ρ*. The black curve shows the results for the asymptotic expressions in Eq. 20 and Eq. 22. Other symbols are as in Fig. 3. Points with error bars show the mean and standard error across 10 runs, each based on 1000 msprime tree sequence simulations. The SMC and SMC^*′*^ curves are obtained by numerically integrating the probability densities derived from Eq. 12 and Eq. 17, respectively. Insets show the error of the analytical expressions that are close to the simulations, with a line at zero added for reference. For high recombination rate *ρ* ≳ *n*^2^, the TMRCA mean and standard deviation scale approximately as 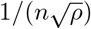. For low recombination rate *ρ* ≪ *n*^2^, the mean and standard deviation approach their single-locus value 2*/*(*n*(*n* −1)). Cumulants under both the SMC and SMC^*′*^ approximations are close to the simulated results, with the SMC^*′*^ being more accurate. The difference between approximations is substantial when the sample size is small and the recombination rate is modest.

In the opposite limit, *ρ* ≫ *n*^2^, we expect recombination to rapidly increase the number of lineages so that the first coalescence occurs quickly compared to a typical coalescence, i.e., at small values of *τ*. This means that we can use the simpler short-time limit for the survival function, which is the same under the SMC and SMC^*′*^ (first line of Eq. 13 or Eq. 18). In this limit, the moments are just Gaussian integrals. The mean TMRCA is approximately:

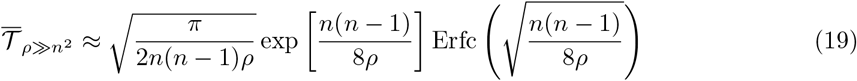

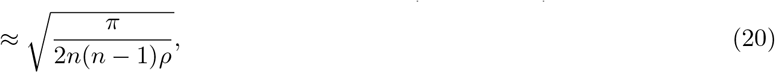

where Erfc is the complementary error function. The variance is approximately:

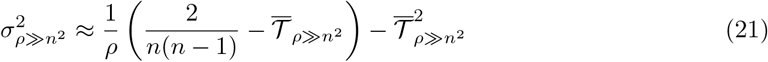

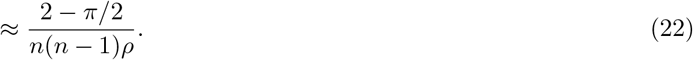

In other words, both the mean and the standard deviation are approximately proportional to 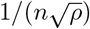 under frequent recombination. In unscaled time, this is 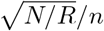, as predicted by our initial intuitive argument, a very different scaling from the single-locus *N/n*^2^. These approximations closely match the numerical results for *ρ* ≫ *n*^2^ (Fig. 4, black line). Notice that in the limit *ρ* ≫ *n*^2^, the SMC and SMC^*′*^ expressions for *F* (first lines of Eq. 13 or Eq. 18) approach our initial heuristic approximation (second line of Eq. 5), so the simple expressions for the mean and variance (Eq. 20 and Eq. 22) are the same as would be obtained directly from the heuristic expression.

### Length of region descended from the most recent common ancestor

We have focused on the time *𝒯* to the most recent common genetic ancestor. One can also consider the length of the region of the genome descended from that ancestor. In the approximately asexual regime *ρ* ≪ *n*^2^, we expect coalescence to occur before recombination, so the entire genomic region under consideration should be descended from the ancestor. In the opposite limit, *ρ* ≫ *n*^2^, recombination will have had time to break the genome up into blocks of length ≈ 1*/ 𝒯* ≪ *ρ*, so we expect that this will be the length of the region descended from that ancestor. More precisely, since we are looking for the overlap between two blocks (one in each of the ancestries that share the MRCA), the expected length should be ≈ 1*/*(2 *𝒯*). There should be some bias towards the region being slightly longer than this, because longer blocks have more partners that they can coalesce with, but simulations show that this is no more than a minor effect when *ρ* is large (Fig. 5).

**Figure 5:**
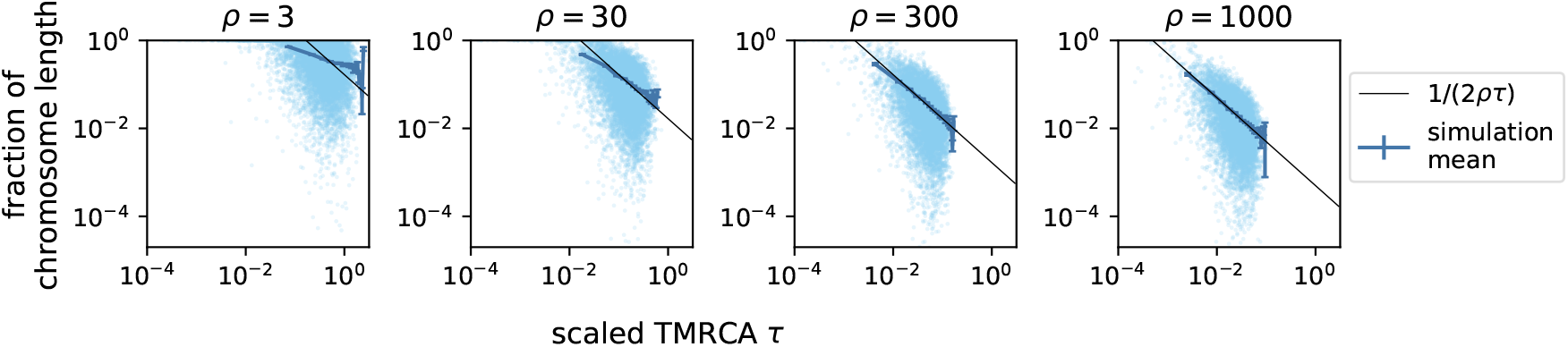
The fraction of the chromosome sharing the most recent genetic common ancestor. Each dot shows the result of one simulation with sample size *n* = 2. When the sampled chromosome has a short map length *ρ* ≲ *n*^2^, the entire chromosome often shares the MRCA. For longer chromosomes with *ρ* ≫ *n*^2^, the mean fraction of the chromosome sharing the MRCA (blue curve) approaches 1*/*(2*ρ*T), inversely proportional to the TMRCA (black line).

## Discussion

In this study, we model the distribution of the time to the most recent common genetic ancestor (TMRCA) of any two individuals in a sample. Using heuristic arguments and the SMC and SMC^*′*^ approximations, we find simple approximate expressions for the distribution and its mean and standard deviation and verify these with full simulations of the coalescent with recombination. We find that there are two possible limiting behaviors, depending on the relative magnitudes of the scaled map length of the sampled genomic region, *ρ* = *NR*, and the sample size *n*. For short regions and large samples, *ρ* ≪ *n*(*n* −)*/*2, coalescence is rapid compared to recombination, and the TMRCA has an exponential distribution independent of *ρ* and occurs when a pair of lineages coalesce across the entire sampled region. The mean TMRCA scales with *N*, the same as the time to the average coalescent event. For long regions and small samples, *ρ* ≫ *n*(*n* − 1)*/*2, the process is quite different. Going back in time, recombination increases the number of ancestral lineages at rate *nR*, increasing the rate of coalescence and creating a TMRCA distribution peaked at a characteristic time 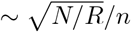 and co rresponding to the coalescence of only a small portion of the sampled region with map length 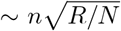. To our knowledge, this is the first time such a scaling has appeared in a simple coalescence time statistic.

To gain some intuition for what these results imply for biologically relevant parameter values, consider an idealized human-like population with a constant size of *N* ∼ 10^4^ and a genome composed of multiple chromosomes, each with map length ∼ 1. If we sample a single chromosome in each of a pair of individuals, then *n* = 2, *R* = 1, and *ρ* = 10^4^ ≫ *n*(*n* − 1)*/*2, so we expect that the most recent genetic common ancestor will have lived 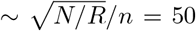 generations ago (∼1000 years ago for human-like generation times), and will be ancestral to only a small portion (∼ 1 cM = 1%) of the chromosome. On the other hand, if we were to sequence only a small *R* = 1 cM region of the chromosome but sample more individuals, say *n* = 100, then we would have *ρ* = 10^2^ ≪ *n*(*n* −1)*/*2, and we would expect the most recent common ancestor to have lived only ∼ 2 generations ago and to be ancestral to the entire sequenced region.

Should one expect to be able to identify the genomic TMRCA and the samples and regions that the MRCA is ancestral to in sequencing data? In the short-region/large-sample limit *ρ* ≪ *n*(*n* − 1)*/*2, it seems likely: the related pair will be closely related over the whole sequenced region, and because the distribution of the TMRCA is exponential, the next most closely related is likely to be substantially more distantly related. But in the more interesting long-region/small-sample limit *ρ*≫ *n*(*n* − 1)*/*2, the TMRCA is likely to occur at a time when the coalescence rate *c* is increasing roughly linearly, meaning that the waiting till the next few coalescent events is likely to be relatively short. Thus it seems likely that in this regime the regions descended from the genomic TMRCA may not stand out from other closely related regions. If extreme outlier regions of high relatedness were found, it would instead suggest that other factors might be playing a role, such as inbreeding or biased sampling of close relatives.

The preceding arguments suggest several ways in which the present analysis could be extended. We considered only a idealized population with constant size, i.e., constant pairwise coalescence rate, but it would also be straightforward to generalize the equations to a time-varying *N* (*t*). In addition, the fact that in the *ρ* ≫ *n*(*n* − 1)*/*2 regime we expect more coalescence events in quick succession after the TMRCA suggests that it could be interesting to calculate further order statistics of the distribution of TMRCAs across the genome. Finally, one could explicitly calculate expectations for observable runs of homozygosity and large blocks of identity-by-descent [Palamara et al., 2012, Carmi et al., 2013, 2014, Browning and Browning, 2015] by taking Laplace transforms of the coalescence time distributions.

## Acknowledgements

The authors thank Stephan Schiffels for suggesting the problem, Peter Ralph for a helpful discussion and for checking our original conjecture via simulation, and Nick Barton, Shai Carmi, and John Wakeley for additional helpful discussions.

## Appendix Discrete-genome simulations

In the main text, we assume that the genomic region under consideration is so long that the chance of two recombination breakpoints coinciding is negligible. Correspondingly, we use a continuous genome in the msprime simulations. Here, we check the effect of relaxing this assumption by simulating a genome with only 100 discrete loci, so that for high recombination rates we expect many independent breakpoints to coincide exactly. The continuous-genome analytical approximations are still good approximations to the TMRCA distribution (Fig. 6) and its mean and standard deviation (Fig. 7). The deviation due to the discrete genome is most apparent for small sample sizes and high recombination, as expected since this is the case in which there is the most recombination before coalescence. But even in this case the mean and standard deviation are accurate to within ≈ 10 − 20%.

**Figure 6:**
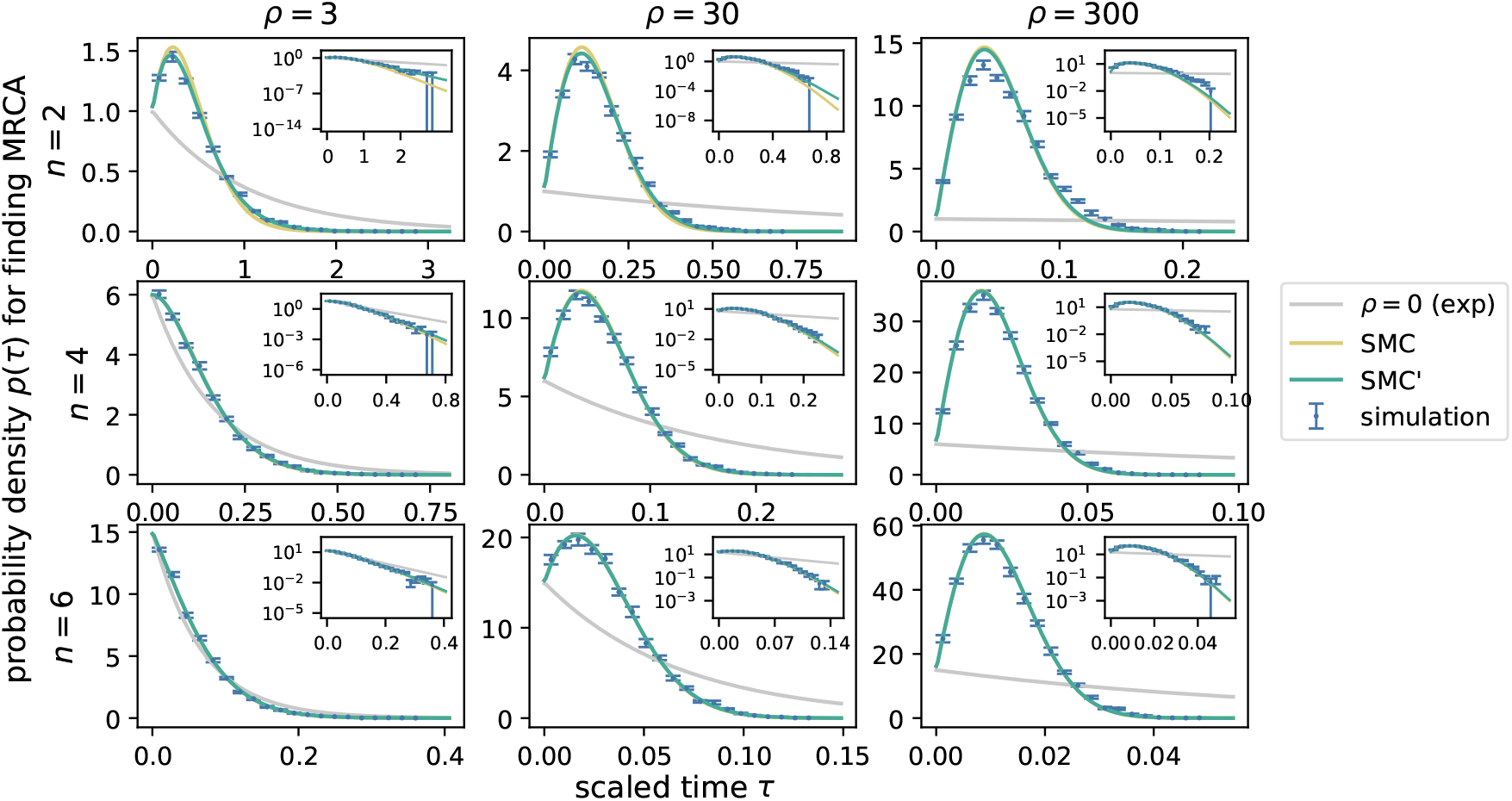
The probability density for the TMRCA. All panels are as in Fig. 3. The only difference is that simulations are for a genomic region comprising 100 discrete loci rather than being continuously divisible by recombination. The only visually apparent difference is for small sample sizes and high recombination rates, where the discrete genome slightly suppresses the peak of the density and lengthens the tail.

**Figure 7:**
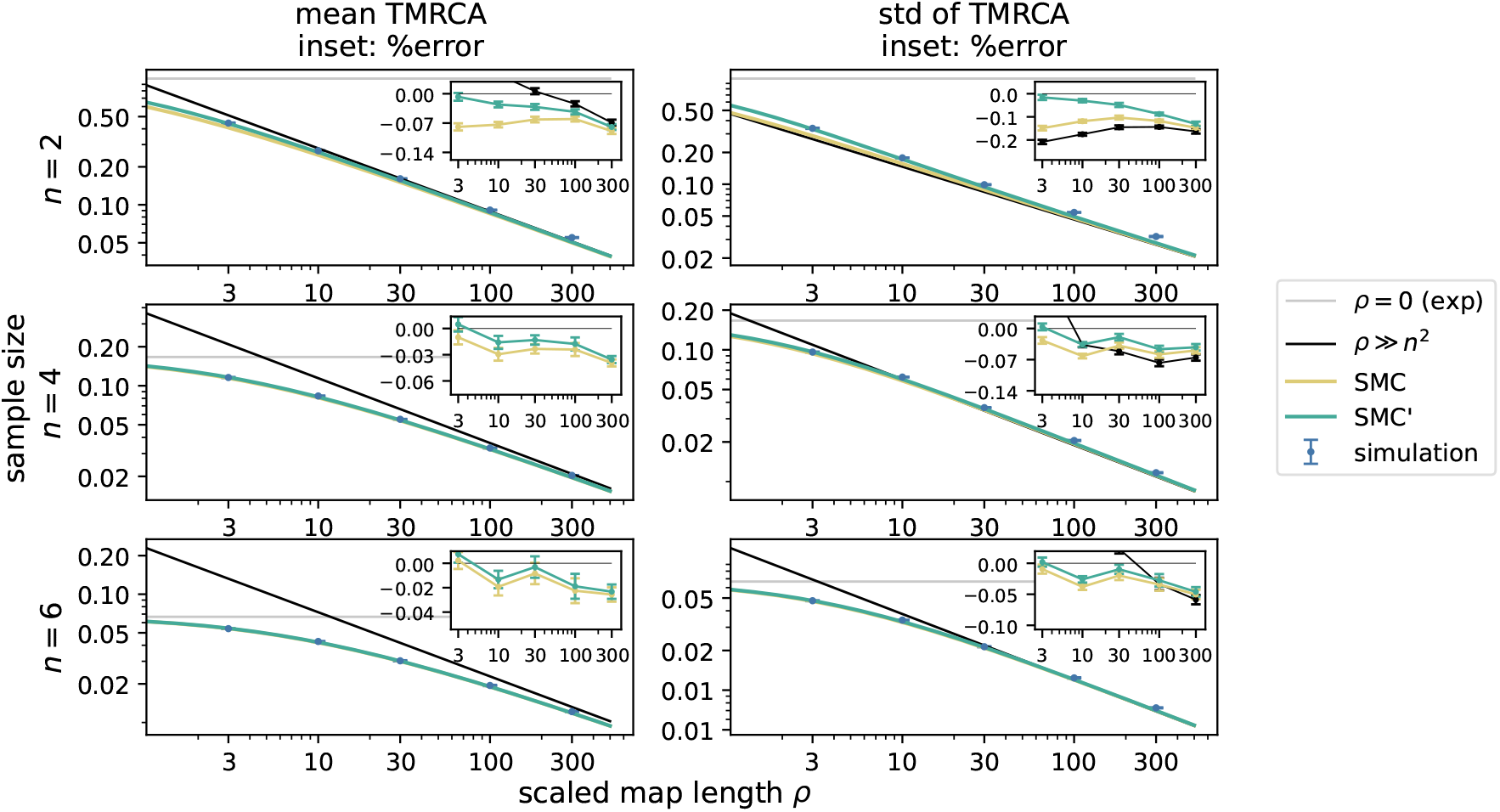
The mean and standard deviation of the TMRCA. All panels are as in Fig. 4. The only difference is that simulations are for a genomic region comprising 100 discrete loci rather than being continuously divisible by recombination. The discrete genome slightly increases both the mean and standard deviation, so that the analytical approximations are underestimates (insets).

## References

Sharon R Browning and Brian L Browning. Accurate non-parametric estimation of recent effective population size from segments of identity by descent. The American Journal of Human Genetics, 97(3):404–418, 2015.

Shai Carmi, Pier Francesco Palamara, Vladimir Vacic, Todd Lencz, Ariel Darvasi, and Itsik Pe’er. The variance of identity-by-descent sharing in the Wright–Fisher model. Genetics, 193(3):911– 928, 2013.

Shai Carmi, Peter R Wilton, John Wakeley, and Itsik Pe’er. A renewal theory approach to IBD sharing. Theoretical population biology, 97:35–48, 2014.

Joseph T Chang. Recent common ancestors of all present-day individuals. Advances in Applied Probability, 31(4):1002–1026, 1999.

Lounès Chikhi, Willy Rodríguez, Cyriel Paris, Marine Ha-Shan, Alexane Jouniaux, Armando Arredondo, Camille Noûs, Simona Grusea, Josué Corujo Inês Lourenço, Simon Boitard, and Olivier Mazet. Extending the IICR to multiple genomes and identification of limitations of some demographic inferential methods. bioRxiv, page 2024.08.16.608273, 2024. doi: 10.1101/2024.08.16.608273.

Bernard Derrida and Bernard Jung-Muller. The genealogical tree of a chromosome. Journal of statistical physics, 94:277–298, 1999.

Robert C Griffiths and Paul Marjoram. Ancestral inference from samples of DNA sequences with recombination. Journal of computational biology, 3(4):479–502, 1996.

Jerome Kelleher, Alison M Etheridge, and Gilean McVean. Efficient coalescent simulation and genealogical analysis for large sample sizes. PLoS computational biology, 12(5):e1004842, 2016.

Jerome Kelleher, Yan Wong, Anthony W Wohns, Chaimaa Fadil, Patrick K Albers, and Gil McVean. Inferring whole-genome histories in large population datasets. Nature genetics, 51(9):1330–1338, 2019.

Leonid Kruglyak. Prospects for whole-genome linkage disequilibrium mapping of common disease genes. Nature genetics, 22(2):139–144, 1999.

Heng Li and Richard Durbin. Inference of human population history from individual whole-genome sequences. Nature, 475(7357):493–496, 2011.

Paul Marjoram and Jeff D Wall. Fast coalescent simulation. BMC Genet, 7(1):1–9, 2006.

Gilean AT McVean and Niall J Cardin. Approximating the coalescent with recombination. Philosophical Transactions of the Royal Society B: Biological Sciences, 360(1459):1387–1393, 2005.

Pier Francesco Palamara, Todd Lencz, Ariel Darvasi, and Itsik Pe’er. Length distributions of identity by descent reveal fine-scale demographic history. The American Journal of Human Genetics, 91(5):809–822, 2012.

Matthew D Rasmussen, Melissa J Hubisz, Ilan Gronau, and Adam Siepel. Genome-wide inference of ancestral recombination graphs. PLoS genetics, 10(5):e1004342, 2014.

Nathan K Schaefer, Beth Shapiro, and Richard E Green. An ancestral recombination graph of human, Neanderthal, and Denisovan genomes. Science advances, 7(29):eabc0776, 2021.

Stephan Schiffels and Richard Durbin. Inferring human population size and separation history from multiple genome sequences. Nature genetics, 46(8):919–925, 2014.

Leo Speidel, Marie Forest, Sinan Shi, and Simon R Myers. A method for genome-wide genealogy estimation for thousands of samples. Nature genetics, 51(9):1321–1329, 2019.

Daniel B Weissman and Oskar Hallatschek. Minimal-assumption inference from population-genomic data. eLife, 6:e24836, 2017. doi: 10.7554/elife.24836.

Peter R Wilton, Shai Carmi, and Asger Hobolth. The SMC′ is a highly accurate approximation to the ancestral recombination graph. Genetics, 200(1):343–355, 2015.

Brian C Zhang, Arjun Biddanda, Árni Freyr Gunnarsson, Fergus Cooper, and Pier Francesco Palamara. Biobank-scale inference of ancestral recombination graphs enables genealogical analysis of complex traits. Nature Genetics, pages 1–9, 2023.

